# Engineering CAR-NK cells targeting CD33 with concomitant extracellular secretion of anti-CD16 antibody revealed superior antitumor effects toward myeloid leukemia

**DOI:** 10.1101/2022.11.14.516308

**Authors:** Rui Zhang, Qingxi Liu, Sa Zhou, Hongpeng He, Mingfeng Zhao, Wenjian Ma

## Abstract

Acute myeloid leukemia (AML) is a common form of acute leukemia and current drugs are overall unsatisfactory. In the present study, we report an immune cell therapy strategy by employing genetically-modified bifunctional CAR-NK cells that combines efficient targeting of AML cells via the CD33 molecule with the concomitant stimulation of NK cell cytotoxicity through the expression and extracellular secretion of anti-CD16 antibody (B16) that binds back to the FC receptor of NK cells. Comparing to CAR-NK cells that target CD33 only, the bifunctional CD33/B16 CAR-NK cells showed superior killing efficiency toward AML cells in vitro, which increased about 4 times based on the number of cells needed to achieve 80% killing activity. In vivo study with xenograft model also revealed effective clearance of leukemic cells and much longer survival - no relapse or death for at least 60 days. In addition, the safety of CAR-NK is not changed with additional expression of B16 as determined by the release of cytokines. These data revealed a promising CAR-NK approach to treat AML patients, which may improve CAR-NK based treatment in general and have potential applications to deal with other tumors as well.

## Introduction

Acute myeloid leukemia (AML) is the most common form of acute leukemia in adults with fairly low survival rates, especially for older individuals [1,2]. Current treatment relies heavily on chemotherapy followed by hematopoietic cell transplantation, which is overall unsatisfactory with a high relapse rate [2]. Novel strategies targeting unique cell surface markers and utilizing the immune system to eliminate leukemic cells have recently become the mainstream of research on this devasting disease [3].

Encouraged by the stellar successes of immunotherapy with chimeric antigen receptor T cells (CAR-T) for patients with hematologic malignancies, CAR-T therapy has received tremendous interest in the treatment of AML [4,5]. Some clinical trials are underway for relapsed/refractory pediatric AML patients (NCT04318678, NCT03971799). However, CAR-T therapy is also facing many challenges such as treatment-related severe toxicity and side effects, including cytokine release syndrome (CRS) and neurotoxicity [6,7]. In addition, a single target antigen as specific and ideal as CD19 or CD22 in B-cells has not been identified for AML [8]. Therefore, non-antigen-specific cellular therapies using natural killer (NK) cells have become an attractive alternative to CAR-based therapy [3].

NK cells are specialized immune cells that can efficiently recognize and lyse malignant cells through the release of cytoplasmic granules, antibody-dependent cellular cytotoxicity (ADCC), or inducing apoptosis [9]. They are key players of the innate immune system to prevent hematologic malignancies and perform immune surveillance functions [10]. Using NK cell as the platform for CAR engineering has the potential to overcome some drawbacks of CAR-T therapy due to their different recognition mechanisms, powerful cytotoxic effects and safety [11,12]. A growing number of preclinical studies indicated that CAR-NK is effective in cancer therapy, particularly in the treatment of hematological malignancies [13,14].

Among the cell surface markers identified so far, CD33 has been found present in 90% of AML blasts [15]. As a myeloid differentiation antigen, CD33 is predominantly involved in myeloid precursors and leukemic stem cells but not in pluripotent hematopoietic stem cells, which makes it an ideal target for antibody-based therapies as well as helps eradicate malignant stem cells that contribute to the occurrence of leukemia [16,17]. A clinical trial of CAR-NK therapy targeting CD33 in patients with relapsed and refractory AML demonstrated that CD33 is a safer target without significant adverse effects [18].

Although targeting CD33 can induce AML cell apoptosis in vitro, the results with unconjugated single CD33 antibodies have so far been disappointing possibly due to the low expression and slow internalization of CD33 which limit ADCC and intracellular accumulation of antibody-delivered payloads [17]. Recently, improved strategies using bispecific antibodies to link surface antigens on tumor cells to receptors of cytotoxic lymphocytes, which redirect immune effector cells to eradicate the leukemia, have shown promising preclinical results [16,19].

CD16 is one of the potential effective conjugates that could work together with CD33 against AML by stimulating the cytotoxicity of NK cells. CD16 is present in the vast majority (>90%) of circulating NK cells [20]. As the only FC receptor that can activate resting NK cells, CD16 plays important roles in mediating NK cells toward eliminating tumor cells through ADCC mechanism [21,22].

In the present study, we developed a CAR-NK system that combines efficient targeting provided by the CD33-CAR molecule toward AML cells with the concomitant induction of stimulatory signals to NK cell itself through the expression and extracellular secretion of anti-CD16 antibody. Our results suggest that it is a promising strategy in the therapy toward AML.

## MATERIALS AND METHODS

### Cell culture and isolation of primary cells

HEK-293T was grown in DMEM (Gibco) supplemented with 10% FCS (Gibco). THP1, U937, MOLM-13 cells were grown in RPMI 1640 medium supplemented with 10% FCS. Peripheral blood (PB) samples from healthy volunteer (n = 3) and relapsed/refractory AML patients (n = 3) were obtained upon approval by the ethics committee of Tianjin the first central hospital.

### Transgene Constructs

Based on the scFv targeting CD33 plasmid (from Creative Biolabs), we constructed a retroviral vector encoding a CD33-specific CAR in combination with the human B16 VHH gene to target the CD16 receptor, which were linked together using 2A sequence peptides derived from foot-and-mouth disease virus and cloned into the PLVX retroviral vector [23]. Self-inactivating baboon envelope-pseudotyped lentiviral vectors (BaEV-LVs) were produced by transient transfection into HEK293T cells using Lipofectamine^™^ 3000 or polyethylenimine (PEI).

### Virus packaging and harvesting

The constructed CD33/B16 CAR-NK plasmid and the virus packaging plasmid (BaEV-gp and SPAX2) were incubated (10ug) with liposome transfection reagent (LipoFiter 3.0) to transfect to 293T cells, and DMEM complete medium was changed 6h after transfection. Cell culture supernatants were collected at 48 h and 72 h, followed by centrifugation at 4,000 rpm for 15min at 4°C and filtration with 0.45 μm filter Membrane, then centrifuge supernatant for 2 hours at 22,000 rpm, and the precipitates were resuspended with fresh DMEM medium to obtain virus concentrate.

### Isolation of NK cells

NK cells were isolated from whole peripheral blood of healthy donors according to published protocols [24]. Briefly, blood samples were mixed with Ficoll lymphocyte separator and centrifuged at 500g for 15min at 4°C. Gently absorbed mononuclear cell layer was resuspended in PBS to centrifuge at 500g for 5min and the resuspended cell pellets were incubated with CD3 Microbeeds (130-050-101, Miltenyi Biotec, Germany) at 4-8°C for 15min. The incubated cells were then sorted with LS magnetic beads (130-042-401, Miltenyi Biotec, Germany), washed with PBS and collected mononuclear cells for reserve. Resuscitated trophoblast cells using NK cell amplification reagent (ZY-NKZ-0104, Hangzhou Zhongying Biology) were then culture in NK cell serum-free medium (supplemented with 5% autologous plasma + 200IU / ml IL-2 + 80U / ml gentamicin) for 3 days as NK cell.

### Transduction of NK cells

Lentiviral particles (MOI=20) and Vectofusin-1 (2.5 μg/ml per well) were mixed in 1ml NK-MACS^®^ medium (ALyS505NK-EX) supplemented with 1% NKMACS^®^ Supplements, 2000 ng/ml IL-1β, 500 IU/ml IL-2 and 10 ng/ml IL-15 (all the above regents were from Miltenyi Biotec). The above transfection mix was added into NK cell culture seeded in six-well plates (3×10^6^/well) and incubated for 24h at 37 °C. Twenty-four hours post-transduction half of the medium was replaced by fresh medium containing 5% human plasma and a combination of IL-2 and IL-15. To detect transfection efficiency, PL protein (RPL-PP2H2, ACRO) was mixed with transfected cell and incubated at 4°C for 30min, washed with PBS and tested by flow cytometer.

### Cytotoxicity detection

To detect the killing efficiency of CAR-NK toward human AML cells, CD33+ cell lines THP1, MOLM-13, and U937 as well as primary cells isolated from M4/M5 PBMCs were used. After incubation with CD33/B16 CAR-NK, CD33 CAR-NK, or control NK cells for 3h and 12h, the cell killing rate were measured by flow cytometry. The killing efficiency were also measured in different cell number ratios between NK cells and AML cell lines at 5:1, 2.5:1, 1:1 and 0.5:1 by flow cytometry.

### Detection of NK cell degranulation and proinflammation factors

NK cell degranulation was determined by the change of CD107a and granzyme. THP1 cells were co-incubated with CD33/B16 CAR-NK, CD33 CAR-NK, or control NK cells at 1:1 ratio for 4 hours, the expression of CD107a and Granzyme were determined by flow cytometry. The cytokine interferon-gamma (IFN-γ) and the growth factor granulocyte–macrophage colony-stimulating factor (GM-CSF) were measured by ELISA.

### In vivo evaluation of CD33/B16 CAR-NK cells in xenografted mice

Female 6-8-week-old NOD/Shi-scid IL-2Rγ(null) (NOG) mice weighing 20±1.6 g (n=36, Vitonlihua Experimental Animal Technology Co., Ltd, Beijing, China) were injected with 3.5×10^6^ THP1 cells expressing luciferase by subcutaneous injection on each side. All animal experiments were approved by the Ethics Committee of Tianjin First Central Hospital (ChiCTR-ONN-16009862). Established tumors were monitored by bioluminescence imaging (BLI). Upon confirmation of engraftment after 7 days, the mice were randomized into 3 groups and treated by tail vein injection of 3.5×10^7^ CD33/B16 CAR-NK or CD33 CAR-NK cells. At 35 days, the CD33/B16 CAR-NK group were treated additionally by tail vein injection of 3.5×10^7^ CD33/B16 CAR-NK cells. Tumor progression were photographed with BLI following intraperitoneal injection with D-luciferin (Goldbio, 150 mg/kg) at 0, 7, 14, 21, 30, 35, 45 and 60 days. All the mice were sacrificed when either experimental or humane endpoints were reached.

## RESULTS

### Engineering CAR-NK cells with bispecific antibodies to simultaneously target AML and stimulate NK cell cytotoxicity

To enhance the specific recognition and killing of NK cells towards AML cells, primary NK cells derived from peripheral blood (PB) were genetically modified by lentiviral transduction to express a second-generation CAR targeting CD33 using a single-chain variable fragment (scFv) derived from the My96 clone of Gemtuzumab ozogamicin (GO), which is an approved antibody-drug conjugate for AML [25]. In order to stimulate the cytotoxicity of NK cells, the human B16 VHH gene, which encodes an antibody that can induce ADCC upon binding to the CD16 receptor on NK cells, was added into the CD33-CAR constructs (Fig. 1A). The CAR and B16 VHH were linked together using 2A sequence peptides derived from foot-and-mouth disease virus to realize the co-translation of the two proteins under the control of the same promoter. (Fig. 1A). Membrane-permeable peptide is used to induce the extracellular secretion of B16 antibody. Thus, generating CAR-NK cells expressing bispecific antibodies (CD33/B16) to simultaneously target tumor cells and trigger NK cell cytotoxicity upon expression.

**Figure 1.**
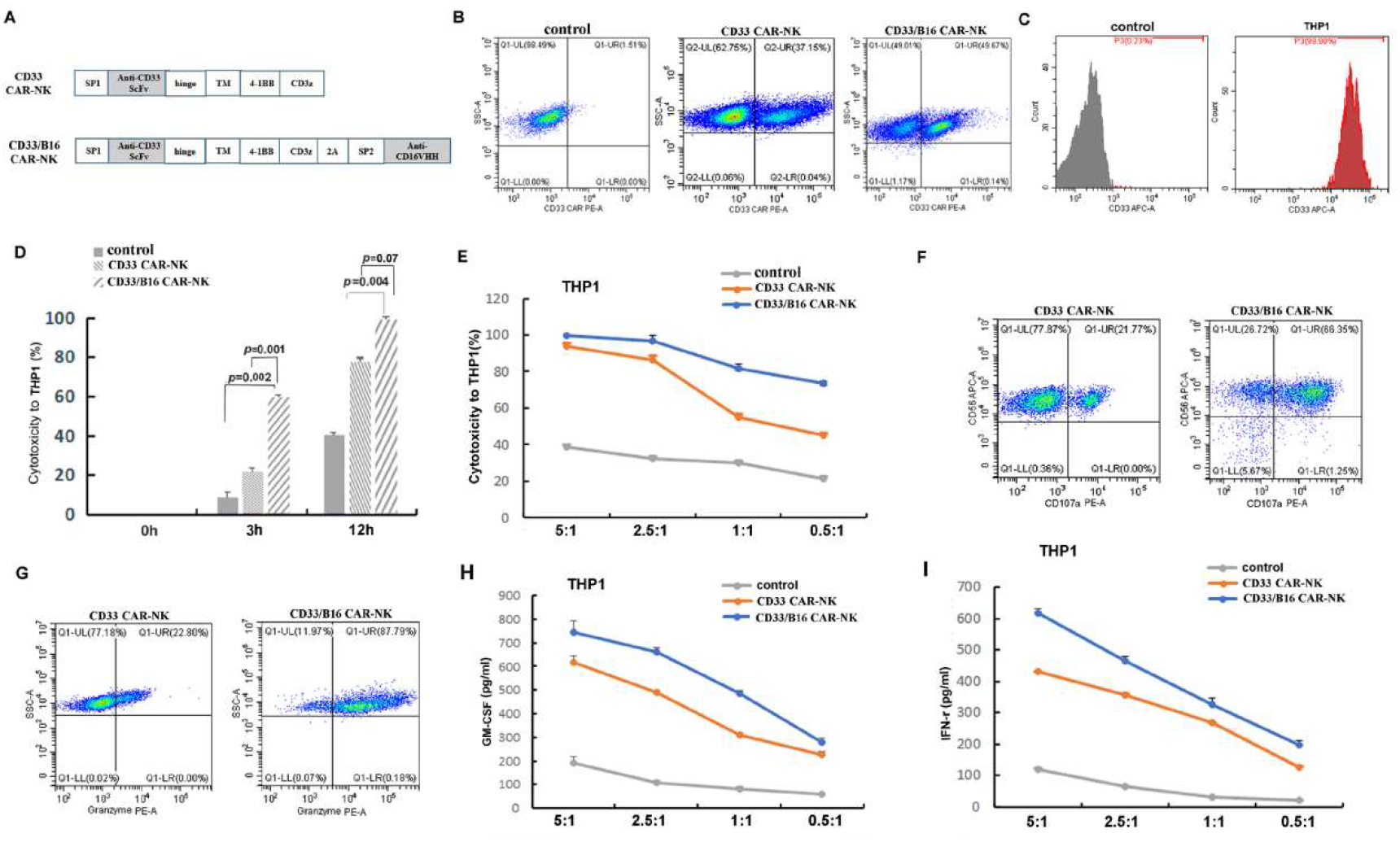
CD33/B16 bifunctional CAR-NK construct and analysis of its cytotoxicity against THP1. A) the design of CD33 CAR-NK and CD33/B16 CAR-NK constructs. B) The expansion of CAR-transduced (CD33-CAR and CD33/B16 CAR) and mock-treated control NK cells in the presence of IL-2 (500 IU/mL) and IL-15 (140 IU/mL). C) THP1 cells are nearly 100% positive on CD33. D) CD33/B16 CAR-NK cells showed much increased killing efficiency toward THP1 cells comparing to CD33 CAR-NK and control NK cells (E:T ratios=1:1). E) the cytotoxicity change at different E:T ratios. F and G) The release of CD107a and Granzyme quantified by flow cytometry. H and I) The secretion of GM-CSF and IFN-γ following CAR-NK treatment of THP1 at different E-T ratios.

### CD33/B16 CAR-NK cells demonstrated potent in vitro activity against THP1

Transgene integration of CD33/B16-CAR and CD33-CAR into NK cells were done by BaEV-LVs with transduction rates between 45% and 60% as determined by FACS (Fig. 1B). In vitro function analysis of the CAR-NK cells was then performed by targeting THP1, a human monocytic cell line derived from AML patients. The expression of CD33 in THP1 was confirmed by FACS which was nearly 100% positive (Fig. 1C). Using a short-term cytotoxicity assay following incubating NK cells and THP1 cells at the E:T-ratio of 1:1, the killing efficiency toward THP1 were determined at 0h, 3h and 12h, respectively. As shown in Fig. 1D, the cytotoxicity of CD33 CAR-NK and CD33/B16 CAR-NK cells were both significantly higher than control NK cells without CAR. In addition, compared to CD33 CAR-NK cells, the killing efficiency of CD33/B16 CAR-NK cells were much more potent, which was about 3 times higher at the time point of 3h post incubation. Further analysis of the cytotoxicity at 6 hours post incubation by changing the ratios of cell number between NK cells and THP cells showed that the cell killing efficiency of B16-CD33 group is always higher than CD33 group unless the E:T-ratio increased to 5:1 (Fig 1E). The smaller the ratio (from 2.5:1, 1:1 to 0.5:1) the more evident the killing efficiency difference between these two groups (Fig 1E), which clearly demonstrated that the presence of B16 gene in the CAR constructs significantly increased the cytotoxicity of CAR-NK cells. To achieve 80% killing activity, it needs about 4 times less CD33/B16 CAR-NK cells than CD33 CAR-NK cells (Fig 1E).

To confirm whether the increased killing efficiency of CD33/B16 CAR-NK cells toward THP1 over that of CD33 CAR-NK cells was due to B16-induced ADCC mechanism, some key marker molecules involved in the activation of NK cell were determined. As shown in Fig 1F and G, a significant increase in mobilization of CD107a (Fig 1F) and granzyme (Fig 1G) were observed upon treatment with CD33/B16 CAR-NK cells comparing to the treatment with CD33 CAR-NK cells. CD107a is a lysosomal-associated membrane protein and its upregulation has been shown correlating with both cytokine secretion and NK cell-mediated lysis of target cells [26]. Granzyme is a serine protease and one of the key components found in the granules of NK cells that is released during ADCC to enter into the cytosol of the target cell to induce apoptosis [27]. These results provided further information on the mechanism underlying the elevated anti leukemic function of CD33/B16-CAR NK cells.

Activated NK cells were known to secrete various pro-inflammatory cytokines and growth factors in dealing with pathological conditions [28]. Therefore, we measured the change of two key markers for this function: the cytokine interferon-gamma (IFN-γ) and the growth factor granulocyte–macrophage colony-stimulating factor (GM-CSF). Comparing to control NK cells, the production of these factors in CD33 CAR-NK and CD33/B16 CAR-NK cells increased ~3-6 times (Fig. 1 H and I), which may account for their sustained killing capacity. Comparing to CD33 CAR-NK, CD33/B16 CAR-NK cells showed further increase of these two factors, but not dramatic.

### CD33/B16 CAR-NK cells demonstrated potent in vitro killing efficiency toward various CD33+ AML cell lines and primary cells from AML patients

To further evaluate the killing efficiency of CD33/B16 CAR-NK cells in comparison to CD33 CAR-NK cells and non-engineered NK cells, in vitro cytotoxicity analysis was performed toward peripheral blood cells of AML patients and some more established AML cell lines. We first determined the expression of CD33 on three primary cell lines obtained from the peripheral blood of relapsed/refractory AML patients (3 cases) and two established human AML cell lines (U937 and MOLM-13). As shown in Fig. 2A, FACS analysis with antibodies against CD33 indicated that U937 and MOLM-13 are mostly CD33+ (>95%). CD33 expression in AML patients varies with one patient showing fully positive (M5 AML patient2) and two patients have a percentage of around 60%. To determine the cell killing efficiency, engineered and control NK cells were incubated with AML primary or established cells for 6 hours at the E:T-ratio of 5:1, 2.5:1, 1:1 and 0.5:1, respectively. As shown in Fig. 2B, the cell killing efficiency of CD33/B16 CAR-NK cells were significantly higher than CD33 CAR-NK cells and control non-engineered NK cells in all AML cell lines tested. Although comparing to established cells lines (MOLM-13 and U937), the lysis efficiency of CAR-NK cells toward AML patient blasts were 10-20% lower, the tendency of improvement of CD33/B16 CAR-NK cells in comparison to CD33 CAR-NK cells is clear, which achieved the same percentage of lysis activity with ~ 4 times less cell numbers (Fig. 2B).

**Figure 2.**
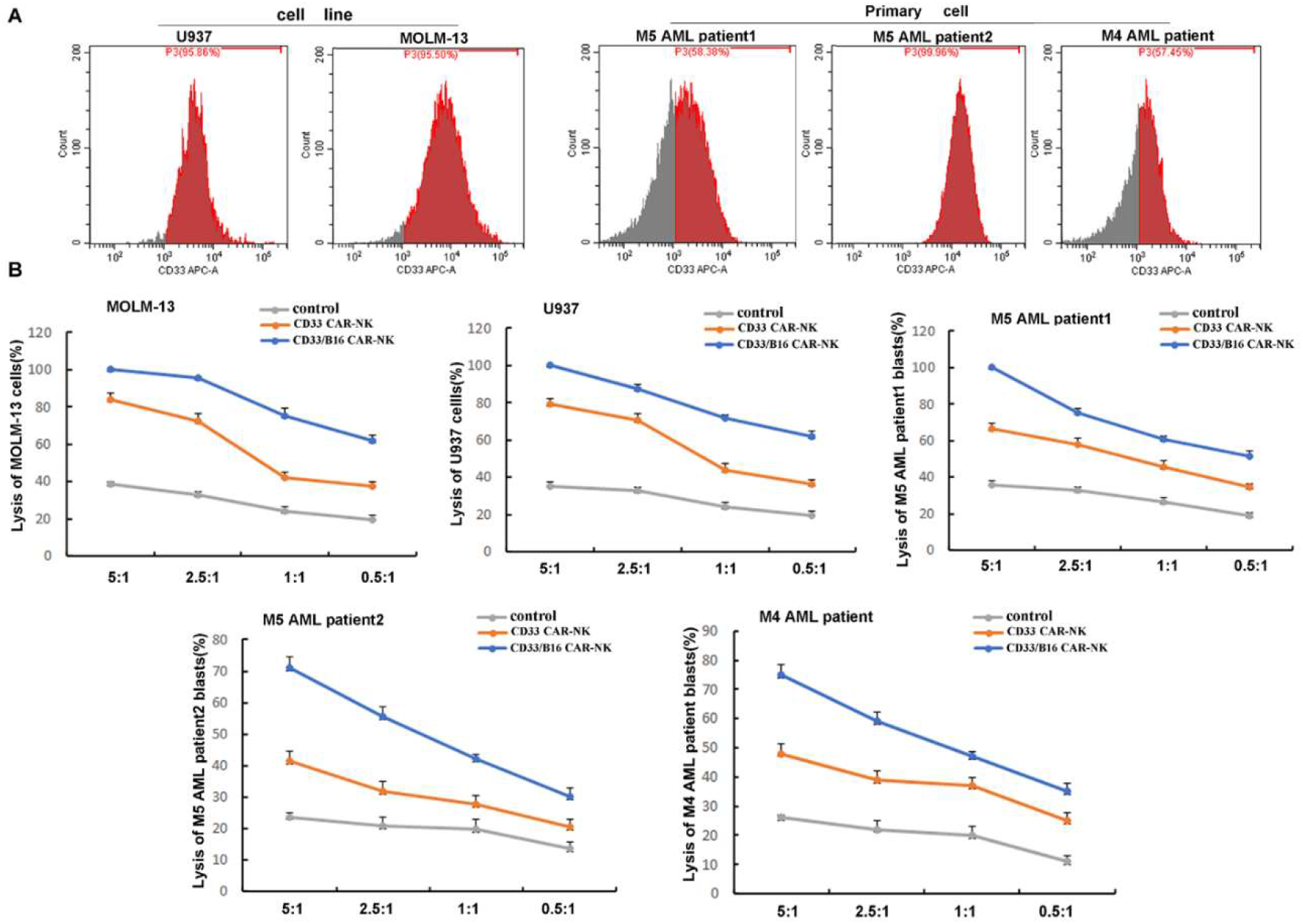
Cytotoxicity analysis of CAR-NK cells against established and primary AML cell lines. A) CD33 expression on established cell lines (U937 and MOLM-13) and primary cells (from M5 and M4 AML patient). B) Comparison of the cell killing efficiency of CAR-NK (CD33 CAR and CD33/B16 CAR-NK) against different AML cells after 6h incubation at E:T ratio of 5:1, 2.5:1, 1:1 and 0.5:1.

### CD33/B16 CAR-NK cells showed effective clearance of leukemic cells in a xenograft model of AML

To evaluate and compare the activity of CD33/B16 CAR-NK and CD33 CAR-NK cells against human primary AML blasts in vivo, they were analyzed in a THP1 xenograft model using NOD/Shi-scid IL-2R γ (NOG) mice (Fig. 3A). NOG mice (male, 6-8 weeks old) received 3×10^6^ THP1 cells (GFP+, Luciferase (Luc)+) intravenously followed by an administration of 3.5×10^7^ CD33/B16 CAR-NK cells or CD33 CAR-NK cells at day 7 post tumor cell injection. Leukemia proliferation was assessed by bioluminescence imaging weekly. Remarkably, a nearly complete eradication of leukemic cells could be observed by day 21 post injection of CD33/B16 CAR-NK cells (Fig. 3A). while a deceleration of leukemic cell growth could be observed in mice treated with CD33 CAR-NK cells at day 14, it didn’t last longer and all mice died by day 35 (Fig. 3A). Leukemic cells reappeared in CD33/B16 CAR-NK treatment group at day 35, we therefore performed a second administration of 3.5×10^7^ CD33/B16 CAR-NK cells at this day (Fig. 3A and B), and mice were all growing well after that for at least 60 days (Fig. 3A). The changes of the average radiance were consistent with the changes of the tumor size (Fig. 3C). The survival time of mice in all the groups demonstrated that mice in CD33/B16 CAR-NK cells had the longest survival with no death (Fig. 3D).

**Figure 3.**
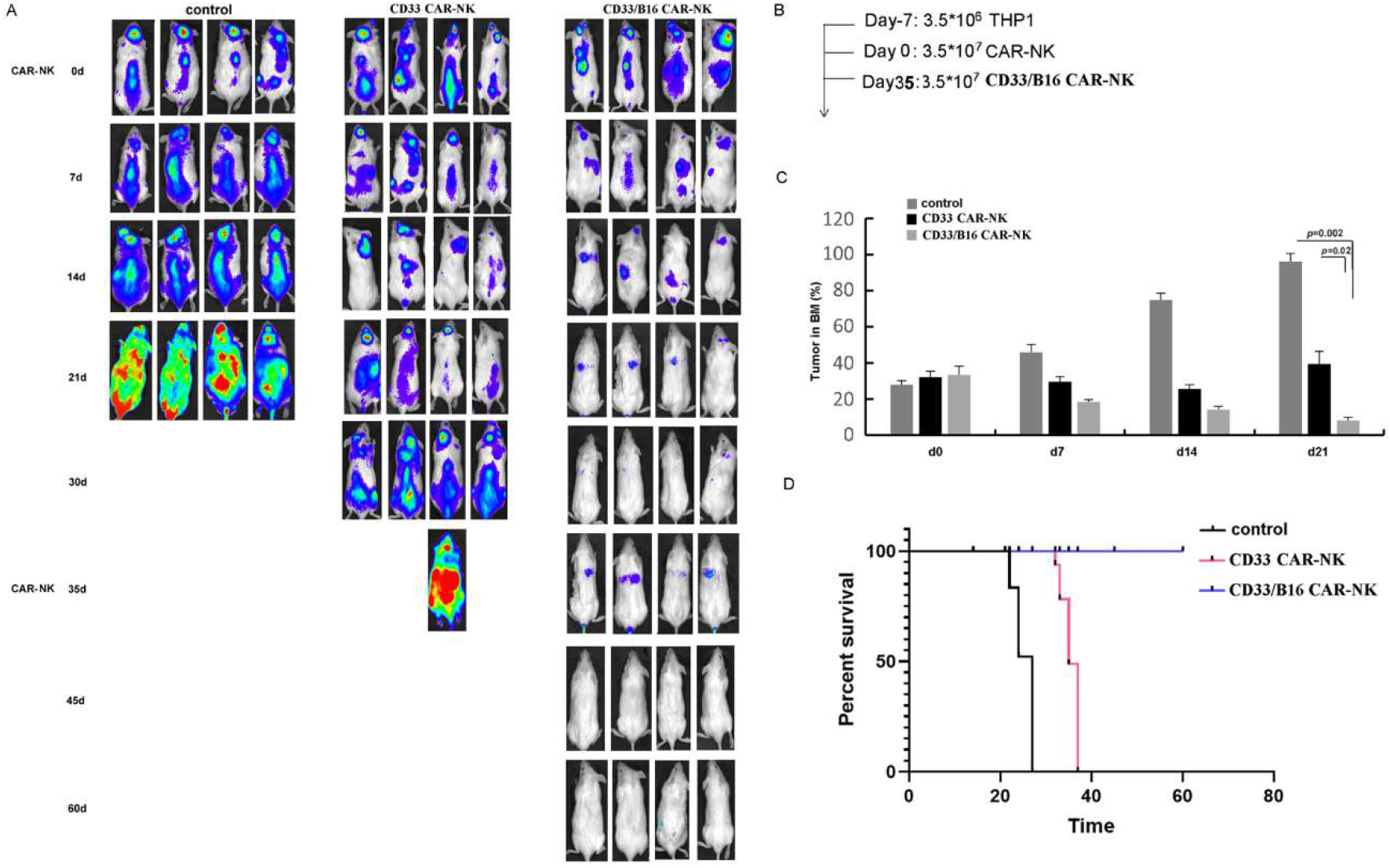
Efficacy analysis of CD33/B16 CAR-NK cells in a xenograft model of AML. A) The formation and progression of tumor were monitored with bioluminescence imaging (BLI) in three groups of mice: control group without any treatment, CD33 CAR-NK group and CD33/B16 CAR-NK group. Day 0 was set when the engraftment was confirmed after injecting the THP1 cells for 7 days (labeled as day-7). B) The time of CAR-NK infusion. C) Change of the tumor size over time. D) The survival rate of three mice groups.

## DISCUSSION

Leukemia patients with co-expression of CD33 usually have poor prognosis, and due to their enormous molecular heterogeneity it is challenging to immune-directed therapies such as CAR-T [29,30]. In the current study, we developed a bifunctional CAR-NK system that simultaneously targeting CD33 and CD16. On the one hand, to bring CAR-NK cells in close proximity of CD33+ leukemic cells. On the other hand, concomitantly stimulating the tumor killing activity of CAR-NK cell itself via the expression of anti-CD16 antibody. Comparing to CAR-NK cells that target CD33 only, CD33/B16 CAR-NK showed superior killing efficiency toward AML cells both in vitro and in a xenograft model.

The cytotoxicity of NK cell is regulated by a repertoire of activating and inhibitory receptors [31,32]. As the only FC receptor that activate resting NK cells, CD16 facilitates the formation of lytic immunological synapse between the NK cell and the aberrant cell (tumor or infected cell), followed by activation of the secretory lysosomes and exocytosis to specifically kill the target cell [22]. CD33 and CD 16 has been used in combination to develop bispecific antibodies, which comprises two antibody fragments that link surface antigens on tumor cells to effector cell receptors of cytotoxic lymphocytes, to trigger antibody-dependent cell-mediated cytotoxicity more efficiently [19,33,34].

In the current study, we adopted the strategy of stimulating NK toxicity through the engagement of CD16 signaling pathway in developing more efficient CAR-NK immunotherapy approach. The expression and extracellular secretion of B16 antibody served as a self-generating stimulator that bind back to the CD16 antigen on the cell surface of NK cells to trigger the cytotoxicity of NK cell through ADCC mechanism. On the other hand, by taking the advantage of CAR approach that drives NK cell directly to tumor cell, it delivers more specific and efficient targeting than bispecific antibodies. Comparing to CAR-NK cells with CD33 only, the cell killing efficiency of CD33/B16 CAR-NK increased about 4 times based on the number of cells needed to achieve 80% killing activity (Fig 1E). Considering that the in vivo amount of NK cell is a limiting factor for its anti-tumor capacity [35], the increased killing efficiency of CD33/B16 CAR-NK provided a promising solution to this obstacle. The therapeutic effectiveness of CD33/B16 CAR-NK was further supported by in vivo studies with the xenograft AML model, which showed striking cure potency in clearing AML cells when compared to CAR-NK cells that target CD33 only (Fig 3).

One of the main advantages of CAR-NK in comparison to CAR-T is that NK cells only secrete a small number of cytokines such as IFN-γ and GM-CSF and do not produce IL-1 and IL-6 that initiate CRS [28]. With the expression of B16 and much increased cytotoxicity, CD33/B16 CAR-NK cells only slightly increased the production of IFN-γ and GM-CSF comparing to CD33 CAR-NK. Therefore, the safety of CAR-NK is not changed by additional expression of anti-CD16 antibody (Fig 1).

Collectively, the current study reported exciting results in improving the killing efficiency of CAR-NK cells toward AML by the concomitant and self-expression of anti-CD16 antibody from genetically-modified NK cell. This approach not only achieves promising anti-leukemia efficacy, but also could potentially reduce the treatment cost by saving the use of additional antibodies, and the strategy may be applicable to solid tumors as well.

## ACKNOWLEDGMENTS

This work was supported by the National Key R&D Program of China (2018YFA0901702), the Shandong Key R&D Program (2019GSF107088), and the National Science Foundation of Shandong (ZR202111220001, ZR2020MC077).

